# A framework for gene mapping in wheat demonstrated using the *Yr7* yellow rust resistance gene

**DOI:** 10.1101/793604

**Authors:** Laura-Jayne Gardiner, Pauline Bansept-Basler, Mohamed El-Soda, Anthony Hall, Donal M. O’Sullivan

**Affiliations:** IBM Research, Warrington, UK; Syngenta, Ferme de Moyencourt, Orgerus, France; Department of Genetics, Faculty of Agriculture, Cairo University, Egypt; Earlham Institute, Norwich, UK; School of Biological Sciences, University of East Anglia, Norwich, UK; School of Agriculture, Policy and Development, University of Reading, Whiteknights, Reading UK

**Keywords:** QTL, association mapping, mapping-by-sequencing, yellow rust, stripe rust, wheat

## Abstract

We used three approaches to map the yellow rust resistance gene *Yr7* and identify associated SNPs in wheat. First, we used a traditional QTL mapping approach using a double haploid (DH) population and mapped *Yr7* to a low-recombination region of chromosome 2B. To fine map the QTL, we then used an association mapping panel. Both populations were SNP array genotyped allowing alignment of QTL and genome-wide association scans based on common segregating SNPs. Analysis of the association panel spanning the QTL interval, narrowed the interval down to a single haplotype block. Finally, we used mapping-by-sequencing of resistant and susceptible DH bulks to identify a candidate gene in the interval showing high homology to a previously suggested *Yr7* candidate and to populate the *Yr7* interval with a higher density of polymorphisms. We highlight the power of combining mapping-by-sequencing, delivering a complete list of gene-based segregating polymorphisms in the interval with the high recombination, low LD precision of the association mapping panel. Our mapping-by-sequencing methodology is applicable to any trait and our results validate the approach in wheat, where with a near complete reference genome sequence, we are able to define a small interval containing the causative gene.

**Highlight:** We show progression from genetic mapping to mapping-by-sequencing and the overlap of defined intervals by each approach culminating with interval refinement and identification of a candidate gene for disease resistance.

## Introduction

Yellow stripe rust (YR) caused by *Puccinia striiformis* is one of the most damaging diseases of wheat with 88% of the world’s wheat production now susceptible to infection (Beddow *et al*., 2015; Ma *et al*., 2015; Singh *et al*., 2016). Therefore, identifying new YR resistance genes and their associated molecular makers is of great interest in breeding programs using marker assisted selection (Gardiner *et al*., 2016; Johnson, 1992). Two major YR resistance genes, *Yr5* and *Yr7*, were identified in wheat and mapped to the long arm of chromosome 2B (Marchal *et al*., 2018; Zhang *et al*., 2009). Intercrossing *Yr5* and *Yr7* near-isogenic lines (NILs) in the genetic background of Avocet S (AVS) revealed distinct segregation patterns for each gene, indicating probable allelism. This hypothesis was confirmed by a recent study which showed that *Yr5* and *Yr7* are distinct paralogues arranged in a complex tandem cluster (Marchal *et al*., 2018). Both are a valuable source of resistance, *Yr5* being effective against the pathogen in most wheat growing regions of the world (McGrann *et al*., 2014) and *Yr7* being effective under various environmental conditions including high temperature (Chen *et al*., 2013).

Genetically dissecting complex responses such as stripe rust resistance is regularly achieved by the analysis of trait-marker associations via QTL and/or genome wide association (GWA) mapping (Agenbag *et al*., 2014; Zegeye *et al*., 2014). While traditional QTL mapping using bi-parental populations is more powerful, GWA mapping can potentially offer higher mapping resolution, as it benefits from historic recombination events and linkage disequilibrium (LD) (Mackay and Powell, 2007). However, GWA mapping can suffer from false positives, leading to false associations, and false negatives, missing true association. Therefore, combining QTL with GWA mapping is increasingly attracting researchers to benefit from both approaches (Brachi *et al*., 2010). Recently, next generation sequencing (NGS) technologies, e.g. mapping-by-sequencing (MbS), has evolved as a new approach to explore trait-marker association with greater resolution via developing well-distributed genic and non-genic SNPs (Gardiner *et al*., 2016; Varshney *et al*., 2014). MbS can be combined with bulked segregant analysis to speed the identification of candidate genes (Takagi *et al*., 2013).

Here, we take advantage of using both bi-parental and GWA mapping populations, phenotyped for severity of infection following inoculation with a pure Pst isolate avirulent on *Yr7*. Both populations were genotyped with the same high-density iSelect SNP array to show the relative power and precision of these population types in locating *Yr7*. We go on to show that using mapping-by-sequencing of resistant and susceptible bulks from the bi-parental population we can identify an overlapping interval to that defined by QTL analysis and GWA. In addition, we can populate the *Yr7* interval with a much higher density of polymorphisms including a series of SNPs from an NBS-LRR gene which would make a logical positional and biological candidate. Performing mapping-by-sequencing of resistant and susceptible bulks using the recent wheat reference genome sequence (RefSeqv1) we show the power of an ordered reference genome. With no prior knowledge of the gene of interest we can refine a genetic interval of 60Mbp containing 589 genes, only 0.55% of the high confidence genes in the genome. Furthermore, with limited information regarding gene function, e.g. in this case our candidate is likely to be an NBS-LRR, we were able to define a candidate gene list of only 10 genes. This methodology is therefore broadly applicable for mapping genes associated with a wide variety of traits. Finally, given that wheat is hexaploid, it was notable that our method is perfectly capable of distinguishing between the homoeologs, identifying a clear peak on chromosome 2B that is absent from 2A and 2D.

## Materials and Methods

### Plant Materials

#### Avalon x Cadenza doubled-haploid population

The population of doubled haploid (DH) individuals, derived from F_1_ progeny of a cross between cultivars ‘Avalon’ and ‘Cadenza’, was developed by Clare Ellerbrook, Liz Sayers and the late Tony Worland (John Innes Centre), as part of a Defra funded project led by ADAS. The parents were originally chosen (to contrast for canopy architecture traits) by Steve Parker (CSL), Tony Worland and Darren Lovell (Rothamsted Research). BS-coded SNP genotypes were imported from supplementary data of (Allen *et al*., 2013).

#### YR-GWAS panel

The main criterion for the membership of the YR-GWAS panel was the existence of at least one year of adult plant resistance data as part of evaluations carried out under the auspices of the National and Recommended List trials and UK Cereal Pathogen Virulence Survey. Based on historic data available at NIAB, 310 varieties, for which well-provenanced seeds could be sourced, were selected to comprise the core wheat YR-GWAS panel. An additional 17 winter varieties were included in the YR-GWAS panel based on their importance in the pedigrees of elite wheat germplasm and the availability of seeds and genotyping information from other projects. The full wheat YR-GWAS panel is therefore composed of 327 wheat varieties, mainly elite UK winter wheat varieties, but other European countries are also represented.

#### Rust phenotyping and the seedling tests

*Puccinia striiformis* (Pst) isolate 08/21 virulent on Avalon and avirulent on Cadenza were used to screen both the AxC DH population and the YR-GWAS panel. Seeds were sown in 96 cell trays, organized in complete randomized block design. Once sown, the trays were placed in a disease-free glasshouse or a growth room prior to inoculation. When the seedlings were between GS11 and GS12, the average height of the seedlings was recorded and plots with average seedling size less than 5 cm, limited seedling number with less than 3, or discoloration were discarded. Following the measurements, the trays were well watered and placed in individual polythene bags. A 1:19 spore: talc mixture was prepared and then 3g of the mixture per tray, equivalent to 0.15g spores and 2.85g of talc, was distributed in an individual glass jar. Using air-blown spore inoculators, each tray was inoculated individually with the contents of a jar. Bags were then sealed and placed in an incubator at 8°C, 24 to 48 hours, in the dark to keep a high humidity level and provide favorable conditions for spore germination. After the incubation, the seedling trays were removed from their bag and placed in a growth room. Two experiments were carried out using the YR-GWAS panel with 2 replications each and one experiment was done with 3 replicates for the AxC DH population. The assessment for the three experiments was achieved 17 days post inoculation. The variety Victor was placed in each tray as a susceptible control and to help choosing the most suitable scoring date. The infection type was assessed on the first leaf of the seedling two times for consistency. Disease assessments followed the 0 to 9 infection type (IF) scoring system described by (McNeal, 1971). Lines with an average IF score < 4 were classified resistant, with IF score 4-6 were interpreted as intermediate response, and with IF>6 were considered susceptible (Table S1).

#### 90k iSelect SNP Genotyping

The wheat 90k Illumina iSelect SNP array (Wang *et al*., 2014) was used to genotype both populations. In case of the AxC DH population, the iSelect mapping data was derived from (Wang *et al*., 2014), whereas genotyping of the YR-GWAS panel revealed 26,015 SNPs.

#### QTL mapping using DH population

Only SNPs on linkage group 2B, i.e. 353 SNPs, which had been scored and were polymorphic in both the AxC and YR panel were used for QTL mapping of seedling test using QTL library in GenStat 17^th^ edition (VSN International, Hemel Hempstead, UK). Default settings of single trait linkage analysis, i.e. 10 cM step size, minimum cofactor proximity of 50 cM, 30 cM minimum separation of selected QTL, and a threshold calculated using Li and Ji option with alpha = 0.05, were used. The QTL was mapped based on interval mapping followed by two rounds of composite interval mapping using cofactors. Finally, to estimate the allelic effect and the explained phenotypic variance, a final QTL model was run.

#### Association mapping using YR-GWAS panel

Kinship matrix and principal components were calculated using Genomic Association and Prediction Integrated Tool (GAPIT) statistical package in R software (Kang *et al*., 2008; Lipka *et al*., 2012). Association mapping was performed using the GAPIT package in R (Lipka *et al*., 2012) using compressed mixed linear model (CMLM) approach that account for population parameters (Zhang *et al*., 2010).

#### Gene enrichment combined with mapping-by-synteny

Resistant and susceptible bulks were created by pooling equal amounts of DNA of 51 individuals with an infection score of >=7 (referred to as the *yr7* bulk) and 54 individuals scoring <=2 (referred to as the *Yr7* bulk). At cost of a potential small loss in genetic resolution, lines with intermediate scores (2.1 – 6.9) were omitted from the bulks to ensure that phenotyping errors did not dilute the allelic discrimination potential of the bulks.

We used the NimbleGen SeqCap EZ in solution custom capture probe set (∼110 Mb) to enrich genomic DNA from the two bulk segregated pools along with the purebred Avalon and Cadenza parental lines. The design and targets of the capture probe set that is used here is as previously reported (Gardiner *et al*., 2014). The final design covered the majority of the wheat genes with each probe intended to enrich all 3 homoeologous gene copies in hexaploid wheat. The samples were paired end sequenced using Illumina HiSeq technology as follows; Genomic DNA was purified using Agencourt AMPure XP beads (Beckman Coulter). Samples were quantified using a Qubit double-stranded DNA Broad Range Assay Kit and Qubit fluorometer (Life Technologies). 2.6μg of genomic DNA, in a total volume of 130μl, was sheared for 3×60s using a Covaris S2 focused-ultrasonicator (duty cycle 10%, intensity 5, 200 cycles per burst using frequency sweeping). The size distribution of the fragmented DNA was assessed on a Bioanalyser High Sensitivity DNA chip (Agilent). 50μl (∼1μg) of sheared DNA was used as input for library preparation. End-Sensitivity DNA chip (Agilent). 50μl (1μg) repair, 3′-adenylation, and adapter ligation were performed according to the Illumina TruSeq DNA Sample Preparation Guide (Revision B, April 2012) without in-line control DNA and without size-selection. Amplification of adapter-ligated DNA (to generate pre-capture libraries), hybridisation to custom wheat NimbleGen sequence capture probes, and washing, recovery and amplification of captured DNA were all carried out according to the NimbleGen Illumina Optimised Plant Sequence Capture User’s Guide (version 2, March 2012), with the exception that purification steps were carried out using Agencourt AMPure XP beads instead of spin columns. Final libraries were quantified by Qubit double-stranded DNA High Sensitivity Assay and the size distribution ascertained on a Bioanalyser High Sensitivity DNA chip. The 4 libraries were then pooled in equimolar amounts based on the aforementioned Qubit and Bioanalyser data. Sequencing was carried out on two lanes of an Illumina HiSeq 2000, using version 3 chemistry, generating 2 x 100bp paired end reads.

The pipeline and algorithm for the processing of this sequencing data is summarized below and in Figure 1; it is adapted from a previously described analysis (Gardiner *et al*., 2014; Gardiner *et al*., 2016). The reference genome that was implemented to map the sequence data was the Chinese Spring hexaploid wheat RefSeqv1 genome assembly. These chromosomal pseudomolecules represent the A, B and D diploid sub-genomes of hexaploid wheat individually resulting in an effectively diploid reference genome of 21 chromosomes (Chapman *et al*., 2015; IWGSC *et al*., 2018). The sequence data was mapped to the pseudomolecules using the mapping software BWA (Li, H. & Durbin, R., 2009). A minimum mapping quality score of 10 was implemented and any non-uniquely mapping reads, duplicate reads or unmapped sequencing reads were removed from the analysis.

**Figure 1.**
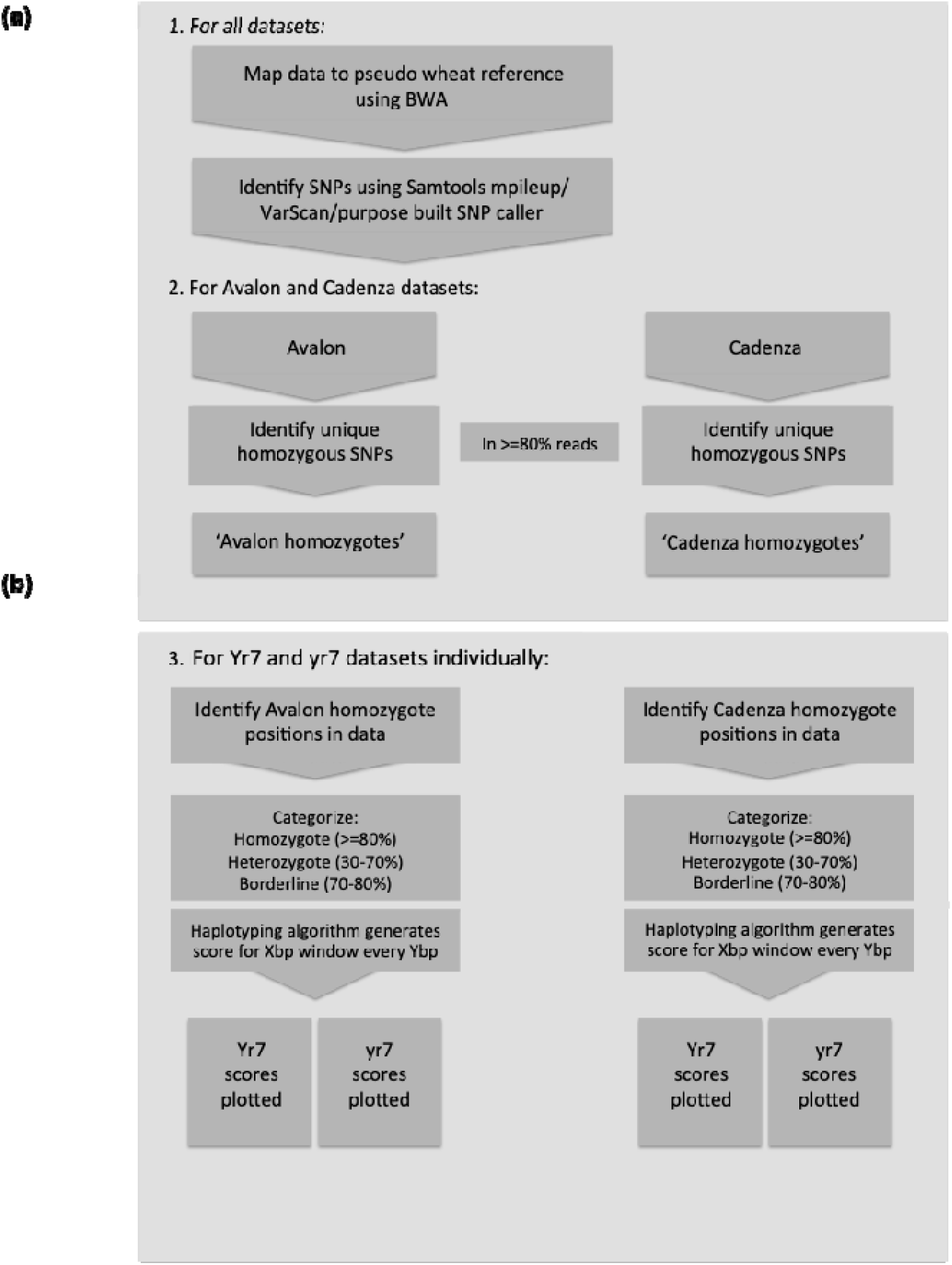
Processing four sets of enriched sequencing data to identify a mapping interval containing the gene controlling the phenotype of interest. **(a)** Standard mapping and SNP calling pipeline to construct “reference genomes” **(b)** Pipeline implementing an algorithm to score regions of interest by prioritizing long homozygous parental haplotypes for the bulk segregant samples to identify the interval of interest. Shown here using Xbp windows at Ybp intervals (user defined) and standard homozygote, heterozygote and borderline SNP definitions for a diploid-values can all be adjusted throughout the analysis as necessary.

SNP calling was carried out on the mapped datasets using Samtools mpileup (Li, H. *et al*., 2009) and then VarScan variant detection (Koboldt, D. *et al*., 2012). SNPs were scored at a minimum quality of 15 and a minimum depth of 10X in Avalon and Cadenza and 20X in the bulk segregant datasets due to their relative average depths of coverage (see results). A maximum depth of 200 was applied and a minimum alternate allele frequency of 10% was used for SNP calls (Table 1). Parental homoeologous homozygous SNPs were called if the SNP allele was observed in 80% or more of the sequencing reads with the reference allele seen in the remainder of the reads. Homozygous SNPs were designated as Avalon- or Cadenza-specific alleles if the homozygous SNP allele was not observed in other parent at that position resulting in 70,895 Avalon and 73,224 Cadenza specific homozygous alleles.

**Table 1.**
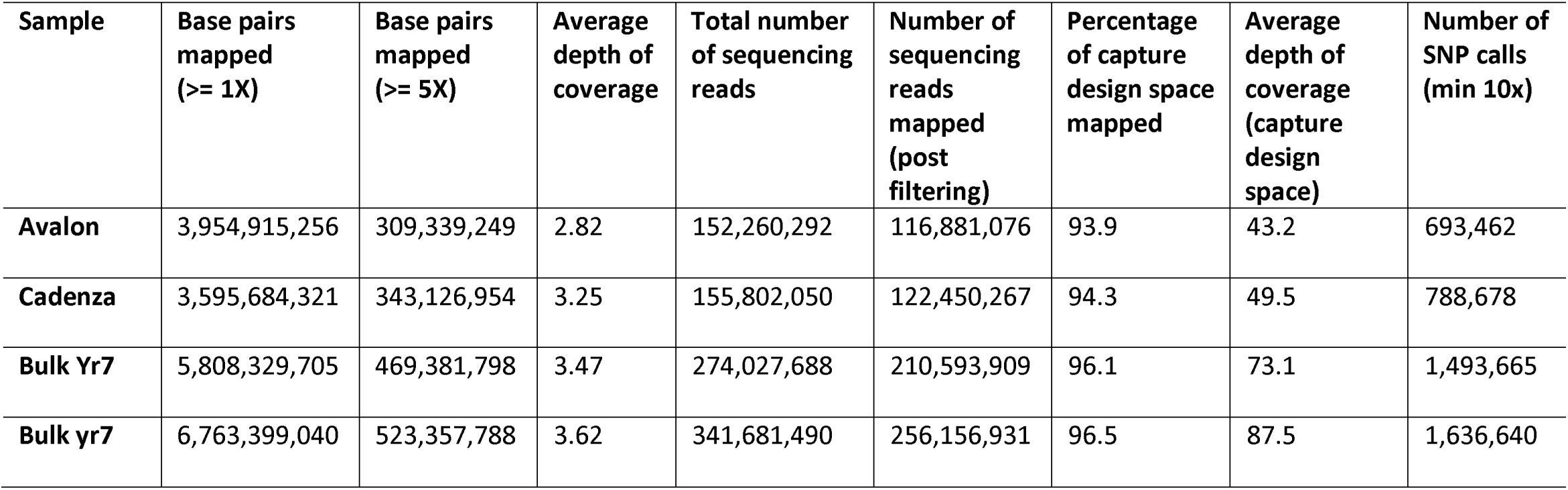
Summary of mapping statistics across the pseudo-chromosome reference. Mapping depth of coverage of the pseudo-chromosome reference and SNP numbers that were identified for the purebred parental lines Avalon and Cadenza plus the bulks

A mapping-by sequencing mutant identification pipeline and homozygous haplotyping algorithm that we presented in a previous study (Gardiner *et al*., 2016; Gardiner *et al*., 2014), were applied here for the analysis of *Yr7* and *yr7* bulks (Figure 1b). Of the Avalon specific homozygous alleles, in regions with a minimum of 20X coverage, 53,658 were seen in the *Yr7* bulk and 60,008 in the *yr7* bulk at the same position, regardless of homozygous or heterozygous status. Similarly of the Cadenza specific homozygous alleles, 51,125 were found in the bulk segregant dataset *Yr7* and 54,946 in *yr7*. These allele positions were then categorized as homozygous, heterozygous or borderline according to the following thresholds; homozygote allele in a minimum of 80% of sequencing reads; heterozygote in 30-70% and borderline in 70-80% (shown in Figure 1b). The scoring algorithm was then implemented to calculate a homozygote score per Xbp window along each pseudo-chromosome at Xbp intervals for these alleles that were found in the bulks and were specific to either the Avalon or Cadenza parents. Window sizes and interval lengths are user defined and adjusted as necessary to suit the dataset and produce the cleanest final plot.

## Results and discussion

### Yr7 maps to a low-recombination region near the centromere of chromosome 2B

Screening of the AxC DH population with Pst isolate 08/21 in a seedling test revealed a bimodal distribution of infection scores, following an approximately 1:1 segregation ratio – the expected inheritance pattern for a single, major gene (Table S2).

A QTL scan for loci conferring resistance to Pst 08/21 showed a single major QTL (-log10(*P*) = 52.6) that explains 70.6% of the variance with a peak at position 110.9 cM on chromosome 2B. Figure 2 shows the CIM QTL trace for chromosome 2B, where it can be noted that the confidence interval spans 29.16 cM. The already large confidence interval, though it spans just 10% of the genetic length of chromosome 2B, contains ∼50% of all 2B AxC SNPs and thus is likely to span an enormous physical interval characterized by low recombination rates, which poses a major challenge for the fine mapping of *Yr7*.

**Figure 2.**
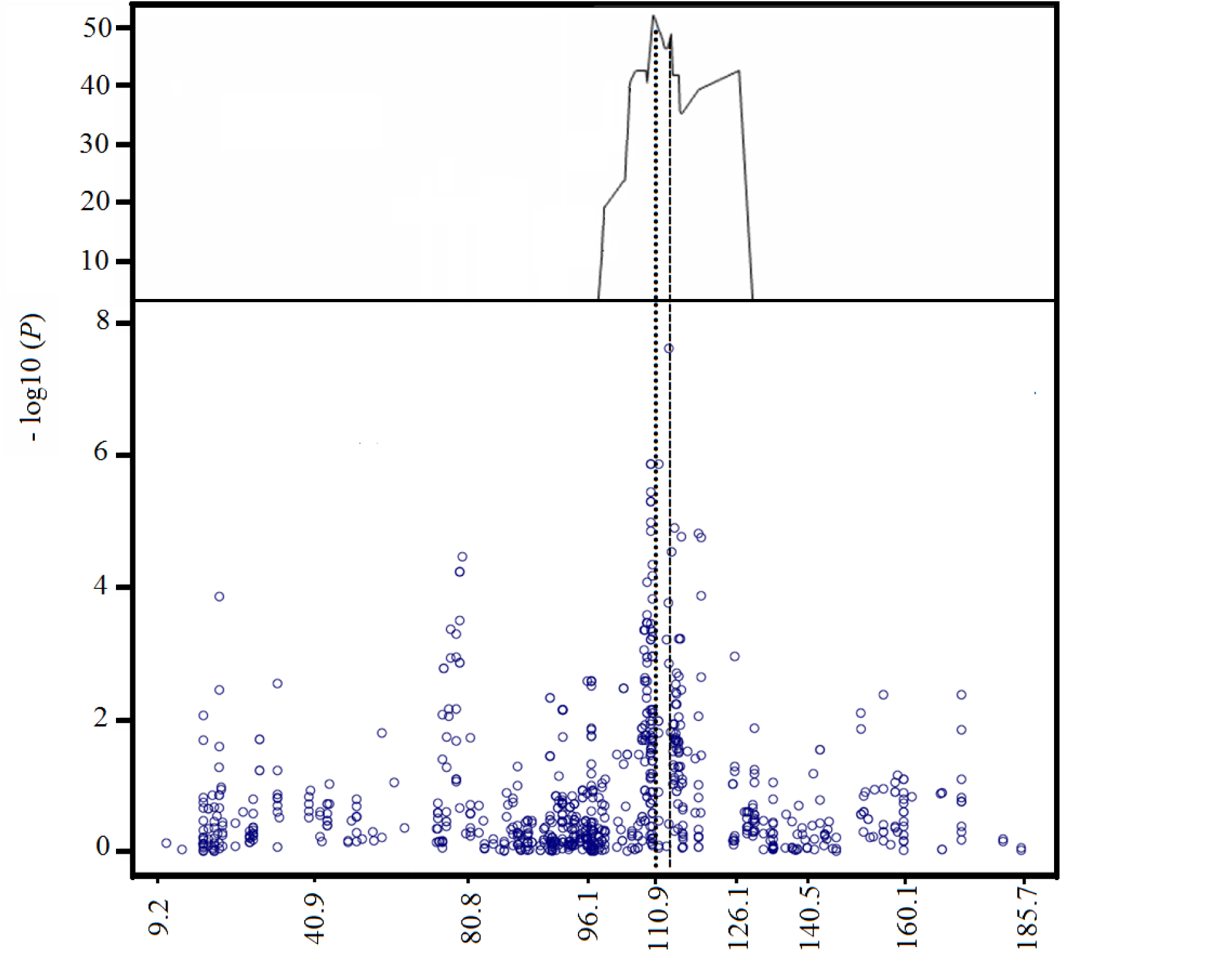
Genetic mapping of the *Yr7* locus. The uppermost plot shows the CIM QTL scan of chromosome 2B for resistance to 08/21 in the AxC DH population. The lower plot shows the GWAS Manhattan plot of chromosome 2B (showing only 353 markers in common with AxC), using AxC map order and distances (cM) to scale the X-axis.

### Genome-wide association mapping reveals historic recombination in the *Yr7* region

In order to gain information on how widespread *Yr7* is in UK winter wheat germplasm, to see the expression of the gene in adult plants grown in the field and to attempt fine-mapping of the *Yr7* gene by another means, the YR Panel of mainly UK winter wheat varieties, which includes both ‘Avalon’ and ‘Cadenza’ as members, was inoculated both in seedling and in adult plant field tests with Pst 08/21. The single significant hit obtained when iSelect SNPs scored in the YR Panel were scanned for association with resistance to Pst 08/21 was located near the centromere on chromosome 2B. The SNP showing the highest probability of association (wsnp_Ex_c10071_16554911) in the GWAS scan is located at 112.9 cM, close to the peak of the QTL interval shown in Figure 2. Closer examination of SNP haplotypes formed by a subset of 47 SNPs informative in both the YR Panel and the AxC population, and located between 107.9 and 119.1cM (see Figure 3) shows that although the *yr7* susceptible haplotype represented by ‘Avalon’ (green shading where ‘Avalon’ allele is present) predominates (251/329) over the resistant ‘Cadenza’ (red shaded) haplotype, several minor haplotypes, some represented by single cultivars, show evidence of recombination between susceptible-like and resistant-like haplotypes. Vertical lines delineating seven recombination ‘blocks’ have been inserted to highlight where multiple distinct and independent haplotypes show blocks of SNPs, presumably in a state of identity-by-descent, have undergone historical recombination. This historic recombination is useful as it allows us both to partly resolve map order, as in the sub-division of SNPs co-segregating in AxC at position 109.24cM into Blocks 2-4, and to tentatively rule out all but Block 6 as potentially carrying the causal gene, since each of the other Blocks have both fully ‘Cadenza’ and fully ‘Avalon’ partial haplotypes in association with susceptibility to Pst 08/21. The notion that graphical genotypes as displayed in Figure 3 have arisen through identity-by-descent in an ancestral recombining population is supported by pedigree analysis which explains 11 out of 17 suspected occurrences of *Yr7* in terms of vertical transmission of haplotypes 25 and 26 through known pedigrees. Haplotype 25 is possessed by varieties **Brock**, Tara and Vault belonging to the same lineage and Haplotype 26 is shared by three separate lineages (1: **Tonic**, Cordiale, Spark, Cadenza; 2: **Ekla**, Vector; 3: **Thatcher**, Tommy). In each case, the founding *Yr7* donor in the lineage is highlighted in bold.

**Figure 3.**
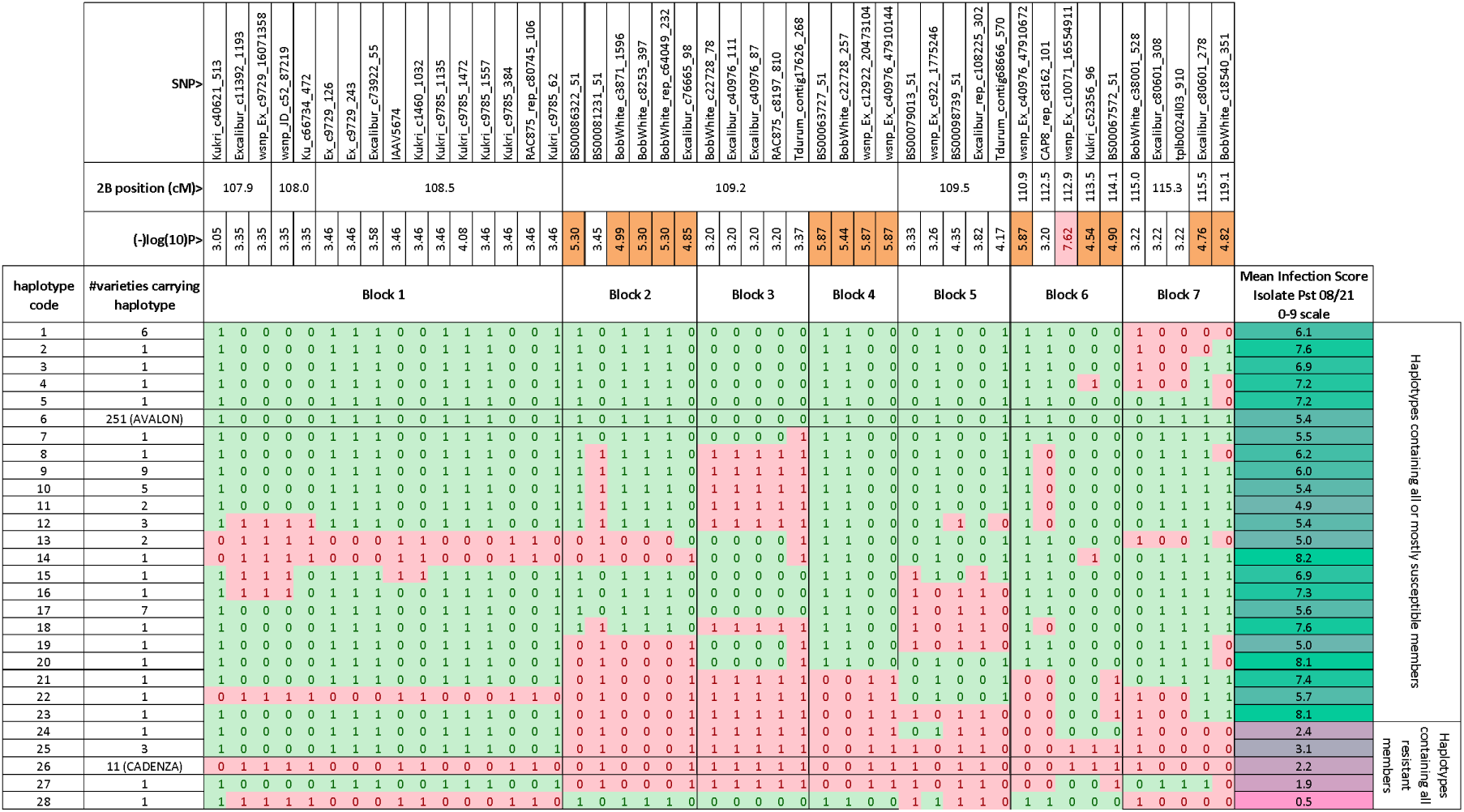
Haplotype structure of the *Yr7* interval in the YR Panel. The 28 distinct haplotypes formed by the 47 SNPs that span the Yr7 confidence interval are displayed in graphical genotype format. For each SNP, the SNP identifier, chromosomal position and P-value for the GWA scan with the Pst 08/21 (Avr-Yr7) isolate are shown. –log10(P)values above 4.5 are highlighted in orange and the most significant value, occurring at 113.8cM in Block 6, is highlighted in pink. The commonest susceptible haplotype (Haplotype 6 which includes ‘Avalon’ and 250 other varieties) is shaded in green and the most common resistant haplotype (Haplotype 26, which includes ‘Cadenza’ and 10 other varieties) is shaded in pink and all other haplotypes are coloured according to which alleles are shared with these reference haplotypes. Mean IF score from inoculation with Pst 08/21 is shown on the right highlighted according to a colour scale that goes from 0 – pink to 9 – green.

### Homozygous haplotyping allows the application of mapping-by-sequencing to hexaploid wheat

After enrichment, the two bulk segregant pools (*Yr7* and *yr7*) were sequenced and mapped along with the purebred Avalon and Cadenza parental lines; over 3.6Gbp of the RefSeqv1 reference was mapped in each of the four datasets with an average depth of coverage of approximately 3.3X. To focus on the targeted regions for enrichment only, reads were aligned to the sequences that were used for the capture probe set design, known as the capture probe design-space. It was observed that ∼95% of this reference was mapped across the four datasets with an average depth of coverage of approximately 45X in Avalon and Cadenza and 80X in the pools *Yr7* and *yr7* (Table 1). Since the capture probe design-space contains one representative copy of each set of three homoeologous wheat genes this translates to ∼15X coverage per wheat sub-genome in Avalon and Cadenza and ∼25X per sub-genome in the pools *Yr7* and *yr7* for the targeted regions.

A mapping-by-sequencing mutant identification pipeline and algorithm that we presented in a previous study (Gardiner *et al*., 2014; Gardiner *et al*., 2016), was here, applied successfully to a new target (Figure 1). Homozygote scores were calculated for a range of windows 100,000-10,000,000bp along each chromosomal pseudomolecule at 10,000bp intervals for the homozygous Avalon and Cadenza specific alleles that could be found in the bulks. The highest signal:noise ratio was obtained when homozygote scores were calculated per 10,000,000bp window at 10,000bp intervals and these scores were plotted for each bulk in relation to both parents in Figure 4.

**Figure 4.**
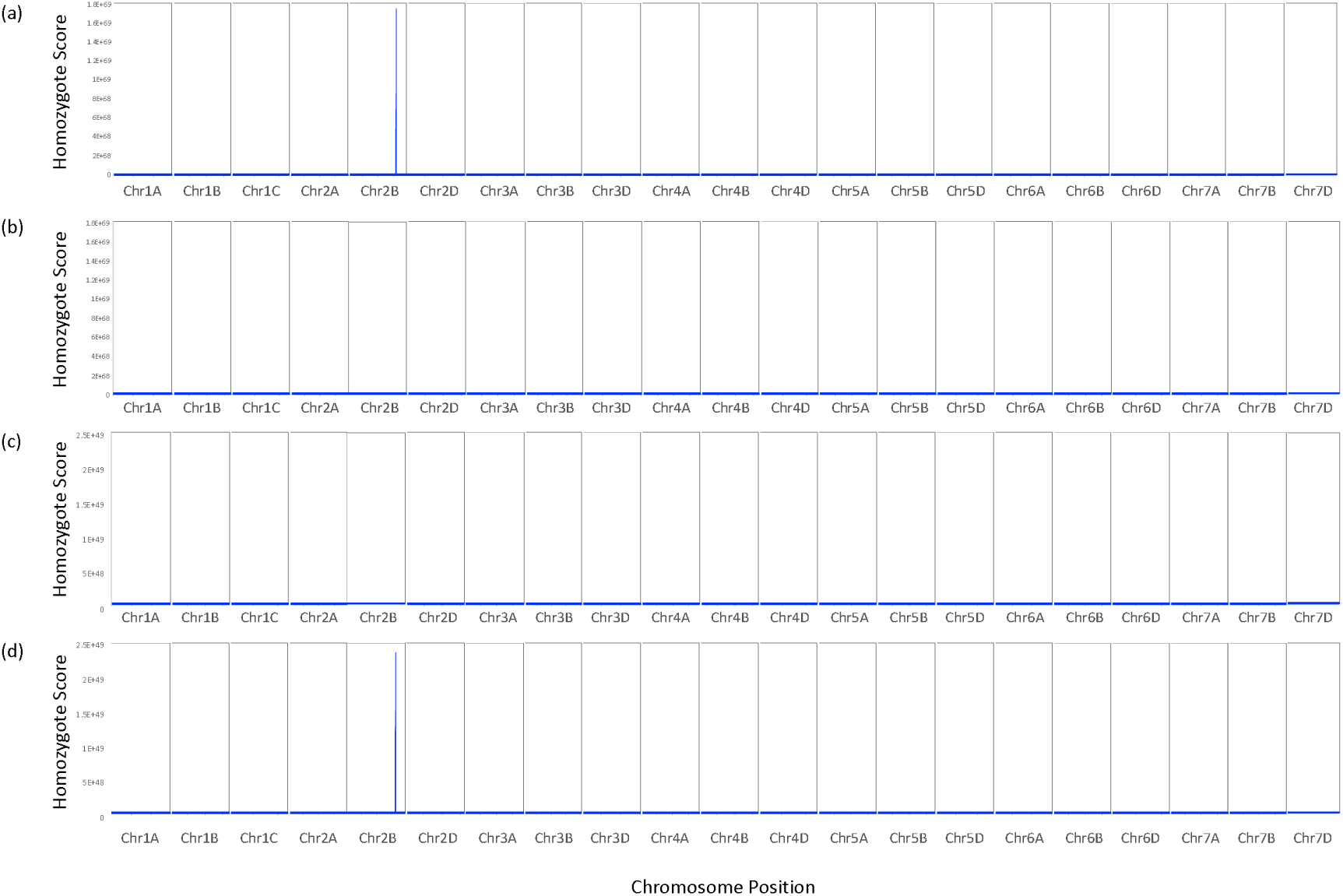
Homozygosity scores calculated for the Yr7 and yr7 bulked datasets along each chromosomal pseudomolecule. Scores calculated per 10,000,000bp window along each chromosome at 10,000bp intervals. **(a)** Scores plotted for ‘Cadenza unique homozygote alleles’ found in the Yr7 bulk segregated dataset. **(b)** Scores plotted for ‘Avalon unique homozygote alleles’ found in the Yr7 bulk segregated dataset. **(c)** Scores plotted for ‘Cadenza unique homozygote alleles’ found in the yr7 bulk segregated dataset. **(d)** Scores plotted for ‘Avalon unique homozygote alleles’ found in the yr7 bulk segregated dataset.

Figure 4 shows a clear peak interval in chromosome 2B that was seen both in the resistant Yr7 pool highlighting conserved homozygosity for the Cadenza parent and in the susceptible yr7 pool highlighting conserved homozygosity for the Avalon parent. The peak interval is shown in greater detail in Figure 5. This is likely to represent our *Yr7* resistance interval of interest as the signal for the *yr7* pool in Cadenza is low, as is the signal for *Yr7* in Avalon (Fig 3b and 3c). The homozygosity scores successfully highlight mirrored ‘Cadenza’-specific and ‘Avalon’-specific peaks in the respective *Yr7* and *yr7* bulks on chromosome 2B with low signal across the rest of the genome. The peak interval encompasses the region from approximately 633,430,001-734,910,001bp on chromosome 2B (homozygosity scores >1e35) which for *Yr7* contains 1,262 ‘Cadenza’-specific SNPs and for *yr7* 823 ‘Avalon’-specific SNPs. Figure 5 highlights multiple individual peak regions that make up the apparently single observed peak in Figure 4; the fragmentation of the signal across disjointed nearby segments is likely a product of the exome capture strategy that was used for sequencing with the multiple peaks representing uneven sequence coverage focused primarily on genic sequence. The highest scoring individual peak regions for *Yr7* Cadenza represent the intervals 633,430,001-663,920,001bp, 681,790,001-693,030,001bp and 702,660,001-721,190,001bp (total of 60,260,000bp with homozygosity scores >1e40). There are 589 high confidence genes observed in these regions.

**Figure 5.**
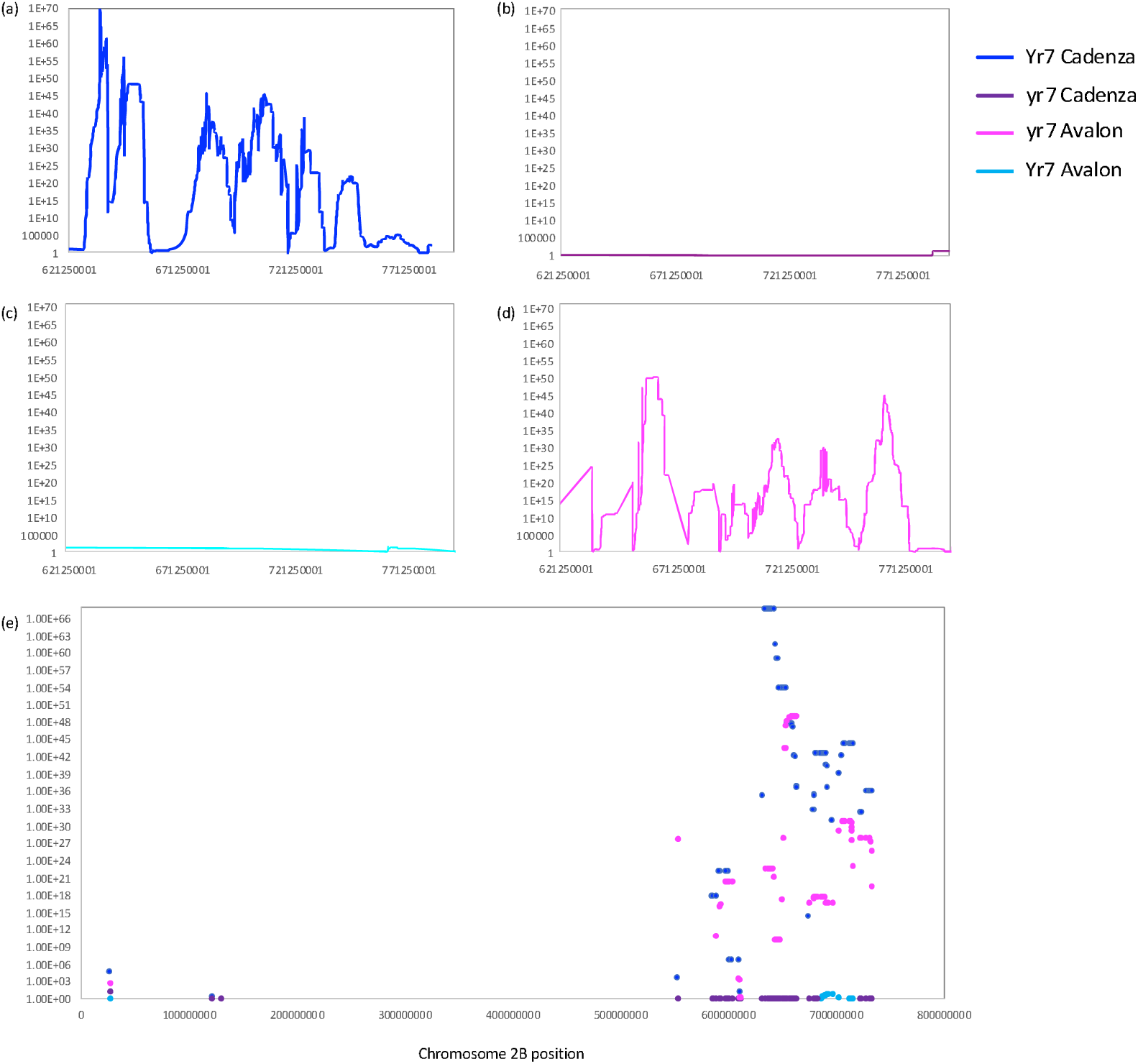
Homozygosity scores calculated for the *Yr7* and *yr7* bulked datasets along chromosome 2B. Scores calculated per 10,000,000bp window along chromosome 2B at 10,000bp intervals. **(a)** Scores plotted for ‘Cadenza unique homozygote alleles’ found in the *Yr7* bulk segregated dataset. **(b)** Scores plotted for ‘Cadenza unique homozygote alleles’ found in the *yr7* bulk segregated dataset. **(c)** Scores plotted for ‘Avalon unique homozygote alleles’ found in the *Yr7* bulk segregated dataset. **(d)** Scores plotted for ‘Avalon unique homozygote alleles’ found in the *yr7* bulk segregated dataset. **(e)** Scores reported here only for windows on chromosome 2B containing the iSelect SNPs that could be mapped to pseudo-chromosome 2B. If multiple windows hit the iSelect SNP, the average score across the windows was calculated. Scores plotted for all datasets **(a-d)** i.e. *Yr7*/*yr7* Avalon and Cadenza specific homozygote scores and datapoints coloured according to the legend.

BLAST analysis was used to infer common features with the sequence (SNP)-based AxC genetic map and the Chinese Spring based RefSeqv1. 296 iSelect SNP sequences from the *Yr7* interval were used in a BLAST search against RefSeqv1 to find their relative positions (E-value >= 1e-5, minimum length 50bp, minimum sequence identity 90%). 277 iSelect SNPs had hits (93.6%) and of these SNPs; 246 (88.8%) were anchored to chromosomal pseudomolecule 2B. The relative homozygosity scores (Cadenza/Avalon specific scores) for these iSelect SNP positions were extracted for each of the *Yr7*/*yr7* datasets and plotted in Figure 5e. Figure 5e shows that the vast majority of *Yr7*-linked iSelect SNP positions on chromosome 2B (216 or 87.8%), fall within the peak region 633,430,001-734,910,001bp that we identified using mapping-by-sequencing. We noted that the 5 SNPs that we previously found in our candidate genetically defined interval ‘Block 6’ (Figure 3), all showed high *Yr7* Cadenza specific (>1E+36) and *yr7* Avalon specific homozygosity scores (>1E+16) but also low Yr7 Avalon specific (<6) and yr7 Cadenza specific homozygosity scores (<1) that are characteristic of the *Yr7* locus. Furthermore, the 5 SNPs were found in the 10.3 Mbp interval 683,029,160-693,314,916bp on chromosome 2B.

### Combining gene enrichment and mapping-by-sequencing for the Yr7 and yr7 DH bulks allows identification of candidate causal SNPs in resistance gene analogues

Once we had established that the combination of; gene enrichment, mapping-by-sequencing and the homozygosity haplotyping algorithm, was specifically and robustly detecting an interval which overlapped the genetically defined interval, the number and nature of *Yr7*-linked polymorphisms could be examined in more detail. The candidate window encompassed 589 high confidence genes, 10 of which are annotated as disease resistance associated and are therefore candidate genes for *Yr7* resistance (Table 2). The 10 genes are mainly clustered together in the region 683,043,955-686,815,417bp (∼4Mbp). In a previous study a candidate *Yr7* gene was suggested and its gene sequence reported (Marchal *et al*., 2018). We used BLAST to align this *Yr7* gene sequence to the Chinese Spring RefSeqv1.1 where its top hit was to the homolog TraesCS2B01G488000 (alignment length 433bp, score 1630, e-value <0.01 and identity 81%). This gene is also seen in the peak interval that was defined here and is one of only 10 of the gene candidates the we report in Table 2. There are 9 homozygous Cadenza specific alleles in the Yr7 bulk in this gene TraesCS2B01G488000 (in a minimum of 70% of the sequencing reads), of which, 4 are predicted to be non-synonymous resulting in codon changes (Table 3). These are potential markers for the *Yr7* gene against a Chinese Spring background. Interestingly, there were no homozygous Avalon specific alleles in the yr7 bulk in this gene, although there was sufficient sequencing coverage in this region to define SNPs. Therefore, it appears that Avalon matches Chinese Spring closely in this region.

**Table 2.** Candidate genes. Annotated disease resistance genes within the candidate intervals of 633,430,001-663,920,001bp, 681,790,001-693,030,001bp and 702,660,001-721,190,001bp on RefSeqv1 chromosome 2B.

| Gene name | Start | End | Annotation |
| --- | --- | --- | --- |
| TraesCS2B01G486100 | 683043955 | 683047878 | NBS-LRR disease resistance protein-like protein |
| TraesCS2B01G486200 | 683053644 | 683059018 | NBS-LRR disease resistance protein-like protein |
| TraesCS2B01G486300 | 683066277 | 683072620 | NBS-LRR disease resistance protein-like protein |
| TraesCS2B01G486400 | 683127582 | 683132111 | NBS-LRR disease resistance protein-like protein |
| TraesCS2B01G486700 | 683159224 | 683163025 | NBS-LRR disease resistance protein-like protein |
| TraesCS2B01G487700 | 683752480 | 683755002 | NBS-LRR disease resistance protein-like protein |
| TraesCS2B01G488000 | 685265721 | 685270904 | NBS-LRR disease resistance protein-like protein |
| TraesCS2B01G488400 | 685741277 | 685746619 | NBS-LRR disease resistance protein-like protein |
| TraesCS2B01G489400 | 686809999 | 686815417 | NBS-LRR disease resistance protein-like protein |
| TraesCS2B01G507100 | 703972543 | 703975392 | NBS-LRR disease resistance protein-like protein |

**Table 3.** Annotation of Cadenza specific alleles that are homozygous in the Yr7 bulk.

| Position | Reference allele | Alternate allele | Concordance | Region | Gene | SNP status | Reference codon | SNP codon | Sample |
| --- | --- | --- | --- | --- | --- | --- | --- | --- | --- |
| 685269082 | A | G | 0.75 | CDS | TraesCS2B01G488000 | Non-Syn | E | G | Yr7 |
| 685269083 | A | C | 0.70 | CDS | TraesCS2B01G488000 | Syn | E | E | Yr7 |
| 685269107 | G | C | 0.72 | CDS | TraesCS2B01G488000 | Syn | L | L | Yr7 |
| 685269174 | G | A | 0.89 | CDS | TraesCS2B01G488000 | Non-Syn | D | A | Yr7 |
| 685269205 | T | C | 0.91 | CDS | TraesCS2B01G488000 | Non-Syn | I | T | Yr7 |
| 685269217 | C | T | 0.82 | CDS | TraesCS2B01G488000 | Non-Syn | A | V | Yr7 |
| 685269221 | G | A | 0.83 | CDS | TraesCS2B01G488000 | Syn | L | L | Yr7 |
| 685269242 | A | G | 0.89 | CDS | TraesCS2B01G488000 | Syn | L | L | Yr7 |
| 685269266 | A | G | 0.91 | CDS | TraesCS2B01G488000 | Syn | L | L | Yr7 |

The 5 SNPs that we previously found in our candidate genetically defined interval ‘Block 6’ that showed the highest –log(P) scores in the GWA scan (Figure 2), were found in the 10Mbp interval 683,029,160-693,314,916bp on chromosome 2B. This interval encompasses our candidate gene homolog TraesCS2B01G488000 (Table 2) that is located at 685,265,721-685,270,904bp. Therefore, gene capture from the same Yr7 and yr7 bulks combined with this mapping-by-sequencing mutant identification pipeline accurately reproduced and refined the results from the genetic mapping analysis to an interval containing 10 genes and ultimately a strong candidate gene. It uses a sliding window scoring algorithm to score regions of interest by prioritizing long homozygous parental haplotypes to smooth out noise in the homozygosity signal.

## Conclusions

### Mapping of *Yr7*

A strong body of evidence links yellow rust resistance present in the cultivars ‘Lee’ and ‘Thatcher’ (used as differential hosts) and their derivatives with the *Yr7* genetic locus. *Yr7* was named by Macer, (1963). Pathotyping studies led to the postulation of *Yr7* in French varieties ‘Camp Remy’ and UK varieties ‘Tonic’, ‘Cadenza’ and ‘Brock’ (Singh *et al*, 2008). *Yr7* has been tentatively mapped by association with QTL in several previous studies. Boukhatem *et al*., (2002) interpreted a QTL (QYR1, gwm501-gwm47) interval on chromosome2B observed in their Camp Remy x Michigan Amber RIL population as *Yr7*. Furthermore, the *Yr7* gene was mapped to the long arm of chromosome 2B (Zhang *et al*., 2009) and recently, a candidate gene validated by isolating several independent susceptible mutants from the ‘Cadenza’ background (Marchal *et al*., 2018). Here we show the progression from genetic mapping to mapping-by-sequencing and the overlap of the defined intervals by each approach culminating with interval refinement and identification of a logical candidate gene using mapping-by-sequencing.

In this work, we are able to re-map *Yr7* with great precision, identifying a set of only 10 gene candidates of which one of the genes was the homolog of *Yr7* in our Chinese Spring reference. We highlight the benefit of combining our mapping-by-sequencing approach with the near complete wheat reference genome sequence (RefSeqv1) in refining our candidate interval. Furthermore, this was possible even though the reference sequence is based on the variety Chinese Spring while our trait was found in the variety Cadenza. The novelty of this study lies in our ability to do this directly in hexaploid wheat from exome capture sequencing data using a mapping-by-sequencing approach that could be applied to any trait, whereas previous approaches use genetic mapping or were able to locate the candidate gene by targeted enrichment of disease resistance genes only and are therefore more specialized in their application (Marchal *et al*., 2018).

### A modified homozygous haplotyping algorithm and mapping-by-sequencing pipeline for hexaploid species

Here, we have taken yellow rust resistant and susceptible bulked doubled-haploid populations, performed gene enrichment and, using mapping-by-sequencing, identified a 60Mbp region on chromosome 2B that contains the *Yr7* locus. The candidate window encompassed 589 high confidence genes, including 10 disease resistance associated genes that included the closest Chinese Spring homolog (TraesCS2B01G488000) of the candidate *Yr7* gene reported previously (Marchal et al., 2018).

This analysis has taken the principles demonstrated by SHOREmap (Schneeberger *et al*., 2009) and applied them to polyploid wheat using a homozygote-scoring algorithm that highlights longer homozygous haplotypes shared between the mutant parental line and the bulked dataset. This extends a proof of concept approach combining genic enrichment and a sliding window mapping-by-synteny analysis using a pseudo-genome. This was firstly carried out in the diploid wheat species *T. monococcum* identifying a region on chromosome 3 that was likely to contain the *Eps-3A*^*m*^ deletion that results in early flowering and later in hexaploid wheat to map the Yr6 yellow rust resistance locus (Gardiner *et al*., 2014; Gardiner *et al*., 2016). Here, this sliding window mapping-by-sequencing analysis, implementing wheat chromosomal pseudomolecules directly, has been successfully applied to define an interval of interest in a polyploid species.

Within the defined candidate gene, we were able to select homozygous alleles at positions that were conserved with the parental lines. This revealed a small subset of 9 Cadenza specific alleles in the Yr7 bulk, of which, 4 are predicted to be non-synonymous and are potential markers for the *Yr7* gene. The identification of a small genetic region containing a candidate gene of interest in a hexaploid demonstrates the power of this analysis and its broad applicability.

### Combining QTL, GWAS and mapping-by-synteny approaches to pinpoint candidate causative polymorphisms

Each of the approaches taken has contributed something to the discovery of potential causative SNPs underlying the *Yr7* locus. The line-by-line SNP genotyping of the biparental population provided an ordered genome scan that treated homoeologues as distinct. The GWAS approach gave further context as to the population history of *Yr7* deployment and the historical recombination evident in the variety panel provided more genetic discrimination than many hundreds more AxC lines could possibly have provided, a fact that was particularly important given the unfavorable physical: genetic distance ratio found in the *Yr7* interval. The nature of this particular YR panel i.e. the fact that it is composed purely of commercial UK winter wheat varieties and important historic founders provides breeders with knowledge of those lineages in which *Yr7* is found and those where it is not, allowing them to pick the most appropriate *Yr7* donor and flanking markers to suit the objectives of the cross and the constraints of their marker assisted selection schemes. Finally, the mapping-by-sequencing defined a list of only 10 genes of which one was the candidate gene itself and also generated additional markers for this gene. We noted that the SNPs that we found in our candidate genetically defined interval closely overlapped the interval defined by mapping-by-sequencing.

### Perspectives for gene cloning in hexaploid wheat

Although the technique of mapping is by now an established method in the genetic toolbox of species which boast a fully assembled genome, polyploid large-genome crop species whose genomes have continued to evade gold standard sequencing and are expensive to re-sequence are still problematic to work with. Trick et al. (2012) showed the application of transcriptome re-sequencing from model bulks in a tetraploid species, which offered one route to the rapid saturation of genetic intervals defined by bulked segregants provided the target genes were expressed to an adequate level. Gardiner et al. (2016) solved the issue of expression variation inherent in the transcriptome resequencing approach by employing exome re-sequencing for gene mapping in hexaploid wheat. In this work, we go a step further by successfully deploying a mapping-by-sequencing approach in a hexaploid using a near complete reference genome sequence. Although the homozygous haplotype scoring map (Figure 4) was initially created blind to knowledge of bulk composition or expected genomic location, it succeeded in remapping the expected interval and in saturating the interval with parent-specific polymorphisms without significant discovery bias and in a manner independent of expression level of the underlying gene. The peak detection sensitivity will depend on the validity of the consensus gene order across all homoeologous genomes and the gene representation in the exome capture. However, evidently this is not an issue for the mapping of *Yr7*.

## Supplementary Data

**Table S1:** Creation of Yr7 and yr7 bulks.

| Gene | R pool cutoff | No. lines in R pool | peak A:B ratio | S pool cutoff | No. lines in S pool | peak B:A ratio |
| --- | --- | --- | --- | --- | --- | --- |
| Yr6 | <=4 | 114 | 0.1875 | >=6 | 69 | 0.0952 |
| Yr6 | **<=2** | **95** | **0.0674** | >=8 | 24 | **0.0000** |
| Yr7 | <=4 | 99 | 0.1034 | >=6 | 73 | 0.0143 |
| Yr7 | <=3 | 89 | 0.0233 | **>=7** | **51** | **0.0000** |
| Yr7 | **<=2** | **54** | **0.0000** | >=8 | 22 | **0.0000** |

R: Resistant lines, S: Susceptible lines

**Table S2.** Segregation of the infection type at seedling stage in AxC population against isolate 08/21.

| Isolate | R | S | Total | Expected<br>Ratio R:S | $\chi^2$ | P-value |
| --- | --- | --- | --- | --- | --- | --- |
| Pst 08/21 | 102 | 99 | 201 | 1:1 | 0.045 | 0.83 |

## Acknowledgments

The authors would like to thank BBSRC for grant BB/J0072607/1 provided to Donal O’Sullivan and the John Oldacre Foundation for Pauline Bansept-Basler’s PhD bursary. This project was supported by the BBSRC via an ERA-CAPS grant BB/N005104/1, BB/N005155/1 (L.-J.G., A.H.) and BBSRC Designing Future Wheat BB/P016855/1 (A.H.).

